# Soil microbiome perturbation impedes growth of *Bouteloua curtipendula* and increases relative abundance of soil microbial pathogens

**DOI:** 10.1101/2024.10.05.616815

**Authors:** Alisiara Hobbs, Daisy Ochoa-Rojas, Christine E. Humphrey, John A. Kyndt, Tyler C. Moore

**Author notes:** Corresponding author: John A. Kyndt.

## Abstract

*Bouteloua curtipendula* (sideoats grama) is a valuable prairie grass for livestock forage, supporting food webs of herbivorous insects, reducing soil erosion, and limiting weed infiltration in urban grasslands. Efficient establishment of *B. curtipendula* in prairie restorations and urban plantings could drastically improve long-term functionality of the space. Soil microbial communities have been linked to plant germination, growth, and drought tolerance in many plant species, however little is known about the factors contributing to *B. curtipendula* germination and early growth. In this study, we used sterilized soil to examine the impact of soil microbes on *B. curtipendula* growth under greenhouse conditions. We found *Bouteloua curtipendula* emergence and growth to be impaired in sterilized soil compared to non-sterilized soil. Using high throughput sequencing of the soil, we found that *B. curtipendula* grown in sterilized soil induced a greater proportion of plant pathogens and fewer nitrifying bacteria when grown in non-sterilized soil. For example, there was a significantly higher proportion of *Acidovorax*, *Cellvibrio*, and *Xanthomonas* which are known to contain plant pathogens, while plant-growth promoting bacteria, like *Rhodopseudomonas,* were significantly higher in the non-sterile conditions. We found that soil sterilization and growth of *B. curtipendula* changed the relative abundance of metabolic subsystem genes in the soil, however, by seven weeks after seeding, *B. curtipendula* transformed the bacterial community of sterile soil such that it was indiscernible from non-sterile soil. In contrast, fungal communities in sterilized soil were still different from non-sterilized soil seven weeks post-seeding. It appears that the bacteria are involved in the initial establishment of beneficial conditions that set the stage for a robust fungal and plant seedling development.

## Introduction

Although classic conservation efforts have focused on expansive nature preserves, recent studies are acknowledging the opportunity for grasslands in cities [1]. Native grassland pockets within cities support insect diversity [2,3], increase pollinator activity [4–6], and increase biological control of pest organisms [7,8]. Urban habitat spaces dominated by regionally-native plants can also enhance the diversity and breeding success of native birds [2,9,10]. In contrast, invasive plant species reduce the capacity for urban plant communities to support bird and insect biodiversity [11,12]. Thus, the capacity for pocket prairies to promote urban biodiversity depends upon the plant community structure.

Because plant composition is an important indicator of urban pocket prairie function, it is important to understand how management efforts could better support diverse native plant communities. One approach to reducing invasive species infiltration to pocket prairies involves a thick planting of native grasses or sedges at the base of forbs to block weed seed access to the soil, shade the soil, and outcompete new seedlings for water and nutrients. This strategy, referred to as “matrix planting” has been shown to be successful in anecdotal reports from landscape designers [13]. Matrix planting relies on a robust clumping or spreading groundcover to fill in the gaps between ornamental forms in a plant community. One commonly utilized matrix plant is *Bouteloua curtipendula*, chosen for its bunching form, drought tolerance, easy establishment from seed, and low growth form [13]. A recent study found broadcast seeding of prairie mixes results in more desirable plant species and less invasive plants compared to more labor-intensive methods (such as planting plugs) [14]. However, strategies that promote rapid and stable establishment of *B. curtipendula* and other matrix plants could therefore increase overall functionality of pocket prairies by contributing to native plant diversity and reducing invasion by exotic plant species.

Soil microbial communities are likely an important, yet understudied, aspect of urban pocket prairie functionality. Soil microbes have intrinsic ecosystem functions which could add to the value of pocket prairies, such as carbon sequestration and reduction of pollutants [15–17]. Soil bacteria mineralize organic nitrogen to more bioavailable forms [18], break down water pollutants passing through soil, and sequester carbon to lessen the impacts of atmospheric carbon dioxide on climate change [19–21]. In fact, urban soil likely provides the same ecosystem services as non-urban soil [17], especially in areas revegetated with locally-native plants [22–23]. These soil microbial processes promoted by urban native plant habitats are likely critical for urban agriculture, human quality of life, and supporting biodiversity within cities.

Soil microbial communities may also be an important factor in determining the rate of establishment of matrix plants in pocket prairies. In order to propagate and establish, plants need to germinate from seeds, uptake water and nutrients from the soil, gather sunlight for photosynthesis (in order to fix carbon dioxide into glucose in the Calvin cycle), and survive stressors which perturb these processes. Soil microbes have long been linked to germination and growth of early seedlings, with both inhibitory and stimulatory effects [24]. Endophytic bacteria have been shown to increase quinoa germination rates by activating immune-related signaling pathways within the seed [25]. A more recent broad scale study used high throughput sequencing to demonstrate a link between soil microbial community and germination across several plant families [26]. Thus, it is possible microbial community processes could promote the germination of matrix plants such as *Bouteloua curtipendula* and thereby increase the success of pocket prairie plantings. Understanding these microbiome-plant relationships in native grasses, such as *B. curtipentula* could potentially open up possibilities for incorporating target strains into seeds, similar to what has been performed in crop plants [27].

Soil bacteria are also able to promote growth of seedlings and mature plants. One mechanism of soil microbe stimulation of plant growth is through enzymes like 1- aminocyclopropane-1-carboxylate (ACC) deaminase, which reduces plant stress hormones such as ethylene [28]. Soil bacterial metabolites also increase water availability [29], which could promote plant establishment during periods of drought. Soil microbial decomposition could also promote plant growth. One study found fungal endophytic symbionts increased plant litter decomposition and subsequent nutrient availability in the soil [30]. Soil bacteria also increase nitrogen and phosphate availability for plants, which could be important for plant growth when these nutrients are limiting [18,19,31]. More recent work has shown the stoichiometric ratio of fungi to bacteria is important for grassland litter decomposition, with low fungi:bacteria ratios corresponding to high nitrogen immobilization by bacteria [32].

Further evidence for the benefits of soil microbes on prairie plant growth comes from work in remnant prairies and prairie restoration. Three herbaceous legume perennials (*Lespedeza capitata, Amorpha canescens, and Dalea purpurea*) had greater plant biomass when inoculated with remnant prairie soil [33]. In addition, remnant prairie inoculation significantly increased the number of leguminous root nodules on *Amorpha canescens* and *Lespedeza capitata* (but not *Dalea purpurea*). However, inoculation did not significantly impact the root colonization into the soil [33]. These results are similar to those showing inoculation of prairie restorations with remnant prairie soil accelerated succession (defined as an increased abundance of late-successional plant species) [34].

The urban setting of pocket prairies makes them unique from prairie remnants, but there is evidence microbes could help plants adapt to these novel conditions. For instance, fitness of *Brassica rapa* during experimental drought was largely linked to microbial communities that rapidly adapted to the new conditions [35]. It is likely that soil microorganisms also increase prairie plant tolerance of elevated heat and drought in urban pocket prairies compared to remnant prairies.

Urban pocket prairies also differ from native remnant prairies in the degree of soil disturbance. In mammalian intestines, disruption of the microbial community with antibiotics increases relative abundance of opportunistic pathogens [36]. Perturbation of soil microbial communities could similarly open niches for less beneficial or even pathogenic bacteria. If so, these disturbances could impede the establishment of prairie plants and reduce the likelihood of urban pocket prairie long-term success. Indeed, soil sterilization reduced the above-ground biomass of *Bouteloua gracilis* [37], but it remains unclear which microbial changes were associated with reduced *B. gracilis* growth.

Although the general role of soil microbes in plant germination, nutrient uptake, growth, and stress tolerance has been described, it remains unclear how soil microbial communities could promote the establishment of matrix plants such as *Bouteloua curtipendula* following acute soil perturbations. In this study, we investigate the role of microbial communities on the growth of the matrix plant *B. curtipendula* by comparing plant growth with or without acute perturbation of the soil microbes (by autoclave sterilization). In addition, we use high throughput sequencing to measure the changes to bacteria and fungi in response to soil perturbation and growth of *B. curtipendula* under controlled conditions. These results could have important implications for the efficient establishment of grasses in urban grassland restorations.

## Methods

### Plant growth measurements

*B. curtipendula* was grown from seed under four conditions; sterilized planted, not sterilized planted, sterilized not planted, and not sterilized, not planted. Standard potting soil and *B. curtipendula* seeds (obtained from Prairie Legacy; SKU BOU01) were used in a 72-cell seed tray. Sterile conditions were created through autoclaving the potting soil for two complete cycles at 132 degrees Celsius for 30 minutes at 15 psi. Soil samples of sterile and not sterilized groups were collected at the time of planting (week 0). Planted conditions received one seed ¼ cm deep in each well. Each condition had 6 replicates, and each replicate was placed in a controlled greenhouse environment with temperature range from 19-25 degrees Celsius, humidity from 60-70%, light levels from 150-425 PAR, and twice daily watering at 5:00 am and 4:00 pm. The height and number of blades were recorded weekly for 7 weeks. Soil samples were collected at weeks 0, 4, and 7. Soil pH was measured by resuspending 500 mg of the soil in 5 ml sterile distilled water and using a pH meter. Roots were separated and dried at week seven to record dry root mass.

### Soil pH measurements and LB agar plating

To measure the soil pH, soil samples were diluted in distilled water and tested with API 5 in 1 test strips. Gross morphological analysis of soil microbiota was performed by plating approximately 1 g of soil on LB nutrient agar plates for one week at room temperature.

### DNA extraction

Soil samples were homogenized, and 250 mg of soil was used for DNA purification using the Qiagen DNeasy PowerSoil Pro Kit according to manufacturer’s protocol (ref: 47014). The extracted DNA was EtOH precipitated for additional cleanup. Samples were resuspended in 4 mM TrisHCl, pH 7.0. The final concentrations ranged from 5.0 ng/ul to 32.7 ng/ul and A260/280 ranged from 1.6 to 2.0. For whole genome-based sequencing we used 100-500ng of each sample and for fungal ITS amplification samples were diluted to 5.0 ng/ul for the initial amplification reaction.

### Next-generation sequencing

Whole genome metagenomics sequencing libraries were prepared using the Illumina Nextera DNA Flex Library Prep kit and were sequenced by an Illumina MiniSeq using 500μl of a 1.8pM library using a High-output cassette. Paired-end (2x150 bp) sequencing generated between 139,594 and 9,540,606 reads (>Q30), and 21 Mbps-1.4 Gbp of data for each of the samples. One replica sample failed to produce sufficient sequencing data (from the sterile soil planted with *B. curtipendula* and harvested at 4 weeks) and was eliminated from the analysis.

Fungal ITS libraries were prepared by using the Fungal Metagenomic Sequencing protocol from Illumina, which uses a pool of 8 forward and 7 reverse amplicon primers that target the fungal ITS1 region between the 18S and 5.8S rRNA genes. They include the ITS1-F and ITS2 amplicon primers from Bellemain et al. [38], that are widely used for fungal barcoding studies. Amplicon primers were synthesized by Sigma Aldrich. Targeted ITS amplicon sequencing was also performed using an Illumina MiniSeq using 500μl of a 1.8pM library using a Mid-output cassette. Duplicate samples were sequenced for each condition. This generated between 517,494 and 1,251,248 reads (>Q30) and between 78.1 and 188.9 Mbp of data per sample.

Both the WGS metagenomic and targeted fungal sequencing datasets were deposited into NCBI Genbank under BioProject PRJNA1163419 and the WGS data are accessible with the following SRA numbers: SRR30751922-SRR30751939 and SRR30754582-SRR30754613. The targeted fungal ITS sequencing files are available with the following SRA numbers: SRR30786940-SRR30786954

### Data analysis

Quality control of the raw reads was performed using FASTQC in BaseSpace (Illumina; version 1.0.0). Adapter sequences were removed and reads with low quality scores (average score < 20) were filtered out using the FASTQ Toolkit within BaseSpace (Illumina, version 2.2.0). Taxonomic classification was analyzed using MG-RAST (version 4.0) [39]. After upload to MG-RAST, data is preprocessed by using SolexaQA [40] to trim low-quality regions from FASTQ data. Potential human sequencing reads were removed using Bowtie [41] (a fast, memory-efficient, short read aligner), and only filtered reads passed into the next stage of the annotation pipeline. MG-RAST uses DEseq for normalization.

Fungal ITS samples were all processed in BaseSpace, where primer and adapter sequences were removed and reads with low quality scores (average score < 20) were filtered out using the FASTQ Toolkit (Illumina, version 2.2.0). The Metagenomics app within BaseSpace (version 1.0.1) was used to perform a taxonomic classification. The UNITE v9 Fungal database (updated November 2022) was used for this analysis.

Default parameters were used for all software unless otherwise noted.

## Results

### Acute soil microbiome perturbation slows growth of B. curtipendula

After one week, *B. curtipendula* sown in sterilized soil visibly germinated in 1/12 samples, while *B. curtipendula* sown in non-sterilized soil visibly germinated in 8/12 samples. By week 2, we observed germination of *B. curtipendula* in 6/12 sterilized soils and 11/12 not sterilized soils (Fig 1A). By week 7, *B. curtipendula* emerged in all samples across all soil types. *B. curtipendula* grown in sterilized soil developed significantly fewer blades of grass compared to *B. curtipendula* grown in non-sterilized soil (P<0.05 by ANOVA) (Fig 1B). By week 7, *B. curtipendula* grown in non-sterilized soil averaged approximately 7.5 blades of grass per sample. In contrast, *B. curtipendula* grown in sterilized soil averaged approximately 2.5 blades per grass at the same time point (Fig 1B). The cumulative blade length of *B. curtipendula* grown in sterilized soil was also significantly less than that of *B. curtipendula* grown in non-sterilized soil (P<0.05 by ANOVA) (Fig 1C), but the difference was mostly observed during the first 4 weeks. By week 6 and 7, we could not detect a significant difference in cumulative *<u>B.</u> <u>curtipendula</u>* grass length between the two conditions (P>0.05 by Tukey post-test for multiple comparisons) (Fig 1C).

**Figure 1.**
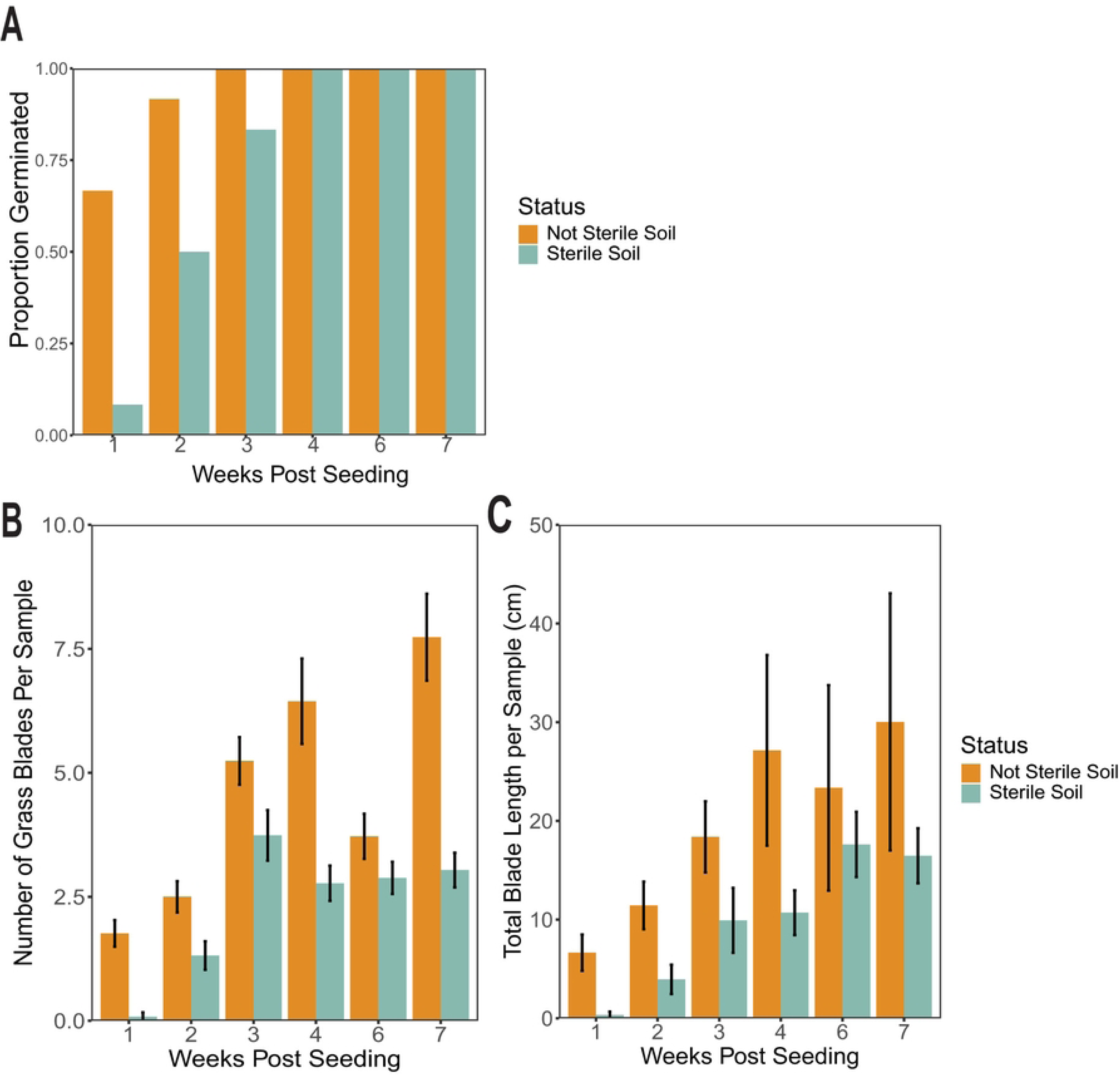
Acute soil perturbation impedes growth of *Bouteloua curtipendula*. Potting soil was sterilized via autoclaving (as described in the materials and methods, “Sterile”) or left unsterilized (“Not Sterile”) and then added to 72 plug trays. *B. curtipendula* was seeded at week 0, and the germination was counted as visible growth of seedlings as a proportion of the total samples (A), and the number of grass blades per plug was counted (B) and the cumulative length of all the grass blades was measured (C). Bars represent means of n=12 samples per group (weeks 1-4) or n=6 samples per group (weeks 6-7) +/- SEM. Differences between means was compared using a one-way ANOVA. (A) Comparison between sterile and not sterile, P<0.001; comparison across weeks, P<0.001; interaction between sterility and weeks, P=0.0138. (B) Comparison between sterile and not sterile, P=0.0006; comparison across weeks, P<0.001, interaction between sterility and weeks, P=0.866.

In order to determine if reduced growth of *B. curtipendula* following soil sterilization was due to changes in soil pH, we measured the pH of soil at time 0 (with or without sterilization), 4 weeks post-planting, and 7 weeks post-planting. All samples had similar pH values (between 8.1 and 8.7), with no notable differences between groups.

### Response of soil microbiome to acute perturbation and B. curtipendula growth

In order to study the soil microbes associated with differential *B. curtipendula* growth, we used whole genome high throughput short read sequencing (HTS) and targeted amplicon HTS of fungus-specific ITS elements. To study the soil bacterial community, we used the WGS output and selected all reads aligned to the domain Bacteria and performed NMDS analysis at the taxonomic level of genus. As expected, we observed a significant effect on the overall soil bacterial community by soil sterilization, however the impact of planting with *B. curtipendula* was significantly different in the sterile versus the non-sterilized soil conditions (P<0.05, Adonis) (Fig 2A). In the non-sterilized samples, soil planted with *B. curtipendula* had a similar overall bacterial community to non-planted soil at both 4 and 7 weeks post-seeding (Fig 2).

**Figure 2.**
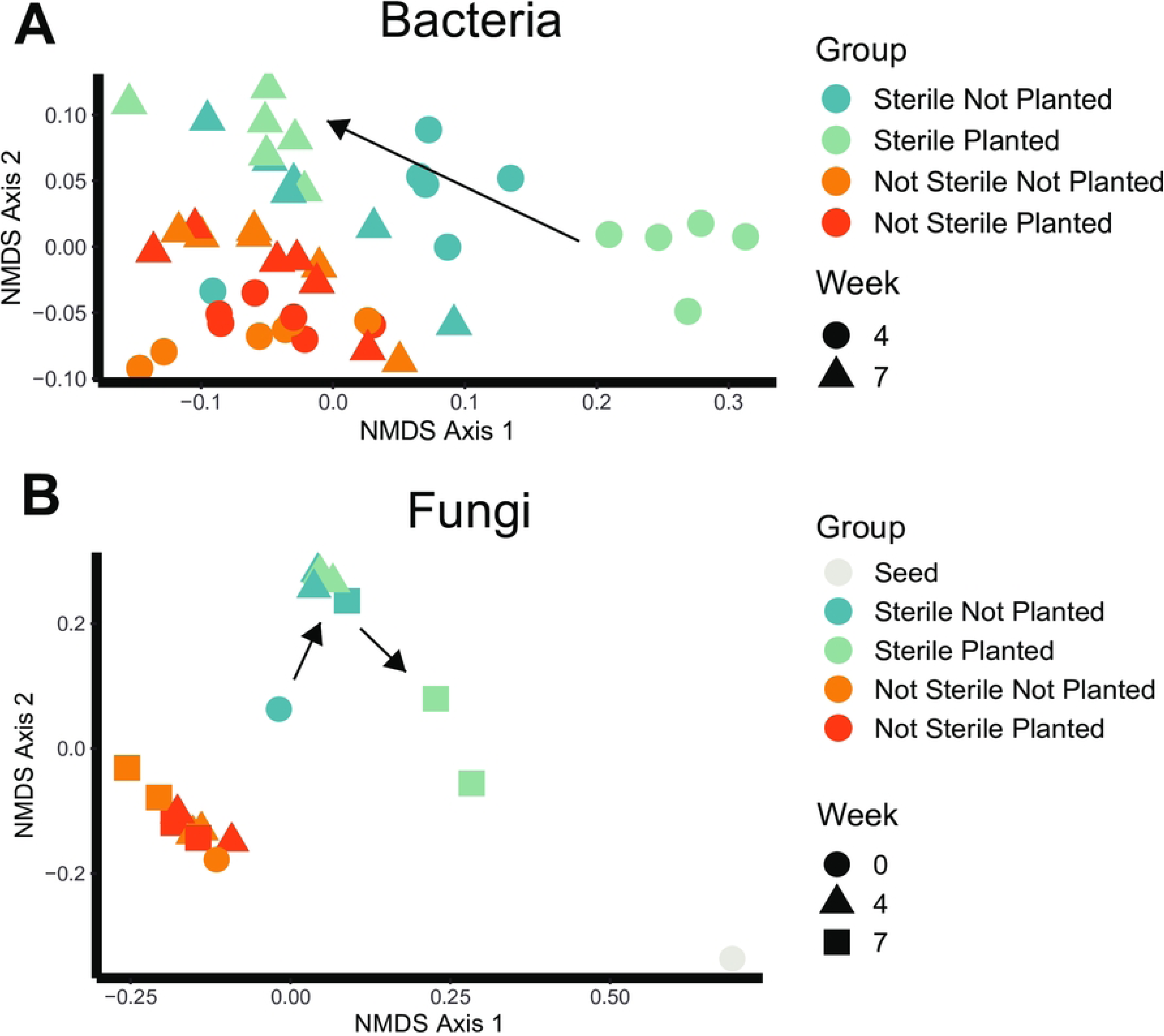
Soil sterilization and planting *Bouteloua curtipendula* shift the soil microbial communities. Potting soil was sterilized via autoclaving (as described in the materials and methods, “Sterile”) or left unsterilized (“Not Sterile”) and then added to 72 plug trays. *B. curtipendula* was seeded at week 0 (“Planted”) or left unseeded as a control (“Not Planted”). At week 4 and seven, soil was collected for DNA purification and illumine sequencing (as described in the materials and methods). The *B. curtipendula* seed (“Seed”) and soil samples immediately after autoclaving (“Week 0”) were taken as a control. Each symbol represents an individual sample. Multi-ordinate variance was compared by Adonis. (A) Comparison between sterile and not sterile, P=0.001; comparison between planted and not planted, P=0.001; comparison across weeks, P=0.001; interaction between sterility and planted/not planted, P=0.011; interaction between sterility and weeks, P=0.001; interaction between weeks and planted/not planted, P=0.001; interaction between sterility, weeks, and planted/not planted, P=0.001. (B) Comparison between sterile and not sterile, P=0.001; comparison between planted and not planted, P=0.002; comparison across weeks, P=0.007; interaction between sterility and planted/not planted, P=0.201; interaction between sterility and weeks, P=0.006; interaction between weeks and planted/not planted, P=0.057; interaction between sterility, weeks, and planted/not planted, P=0.063. Arrows indicate changes from week 4 to week 7 in the sterilized soil microbial community following planting with *B. curtipendula*.

However, in the sterilized samples, soil planted with *B. curtipendula* had a unique bacterial community compared to unplanted soil at 4 weeks post-seeding. By 7 weeks post-seeding into the sterilized soil, the bacterial community in both planted and unplanted soils converged toward the bacterial community of non-sterilized soils (Fig 2A). These results indicate a transient disruption to the soil bacterial community resulting from planting *B. curtipendula* into sterilized soil, which did not occur in the non- sterilized soil.

We also observed a significant interaction between soil sterilization and *B. curtipendula* planting on the soil fungal community (P<0.05, Adonis). In contrast to what we observed in bacteria, fungal communities in sterilized soil were similar between planted and unplanted samples at week 4 (Fig 2A). Instead, planting *B. curtipendula* into sterilized soil induced a unique fungal community compared to unplanted sterilized soil which can be observed as a significantly different mycobiome at week 7 (Fig 2B). Interestingly, fungal communities in sterilized soil planted with *B. curtipendula* began to more closely resemble the seed fungal community by week 7, suggesting seed-origin fungi were better able to colonize sterilized soil (Fig 2B).

### Functional diversity changes in response to soil microbiome perturbation and planting with B. curtipendula

At week 0, sterilized soil had drastically less bacterial diversity and richness than nonsterile soil (Fig 3A, 3C). Bacterial diversity and richness of the seed before planting it (week 0) was intermediate between sterile and non-sterile soil (Fig 3A, 3C). We observed slightly more bacterial diversity and richness in sterilized soil planted with *B. curtipendula* compared to unplanted soils at 4 weeks post-seeding (Fig 3A, 3C).

**Figure 3.**
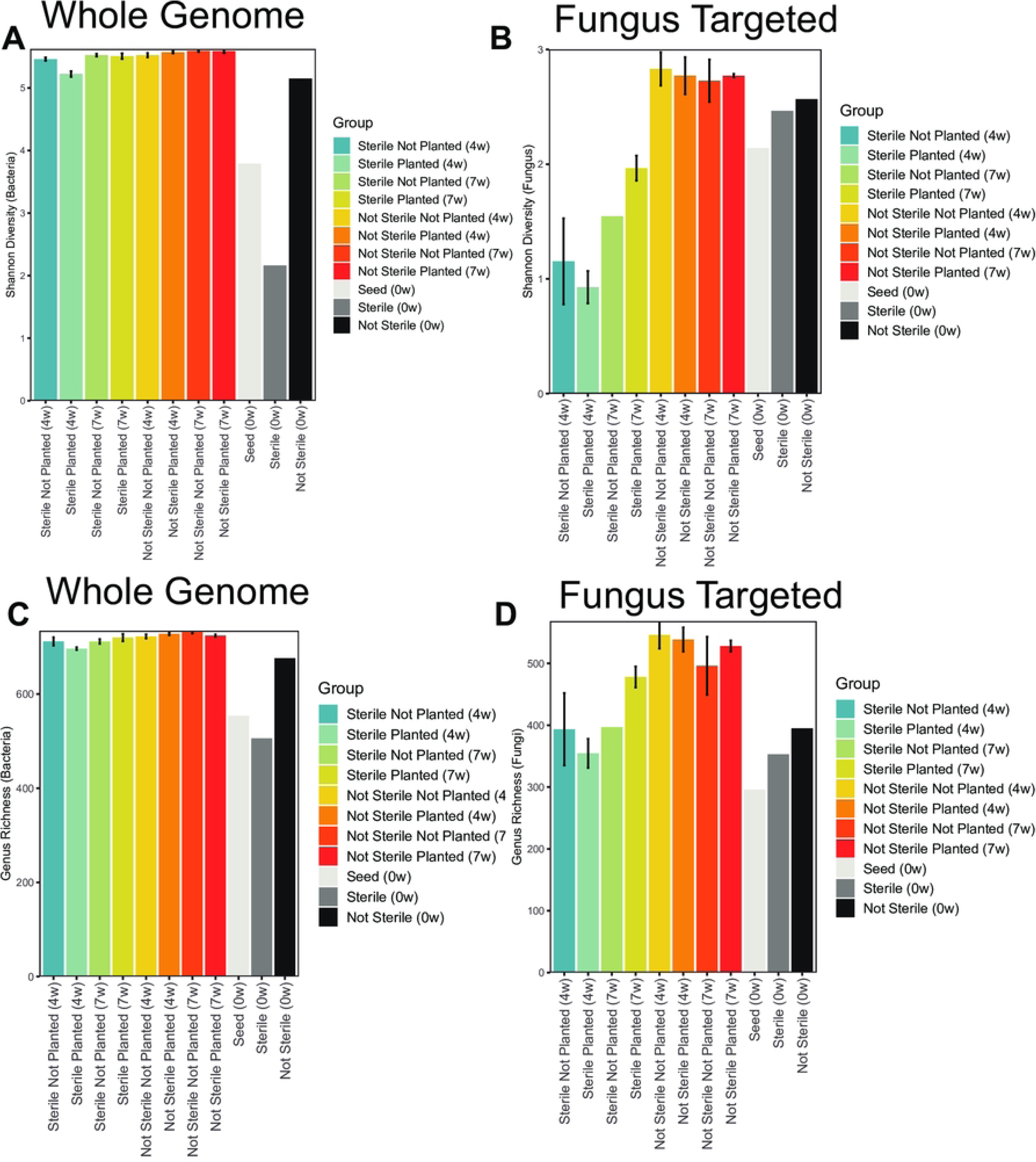
Soil microbial diversity and richness changes in response to soil sterilization and *B. curtipendula* growth. Potting soil was sterilized via autoclaving (as described in the materials and methods, “Sterile”) or left unsterilized (“Not Sterile”) and then added to 72 plug trays. *B. curtipendula* was seeded at week 0 (“Planted”) or left unseeded as a control (“Not Planted”). At week 4 and seven, soil was collected for DNA purification and Illumina sequencing. Bars represent means +/- SEM of diversity (A, B) or richness (C, D) within whole genome sequencing (A, C) or fungus ITS-targeted sequencing (B, D), n=6 samples per group (week 0 and seed, n=1).

However, bacterial richness and diversity were equal across all conditions by 7 weeks post-seeding (Fig 3A, 3C). In contrast, we did not observe a decrease in fungal diversity immediately following sterilization (week 0) (Fig 3B). However, sterilized soil at 4 weeks showed significantly less fungal diversity than not sterilized soil. Although fungal diversity of the sterilized soil increased from 4 weeks to 7 weeks, diversity was not fully restored to levels observed in not sterilized soil. Of note, we observed more fungal diversity and richness at 7 weeks in sterilized soil planted with *B. curtipendula* compared to unplanted soil.

Figure 4 shows the metabolic pathway groupings (level 1 subsystem) from whole genome sequencing. Carbohydrates, clustering-based subsystems, amino acids and derivatives, and protein metabolism were the most abundant across all groups. Among the sterilized soil at week 0, carbohydrates and dormancy and sporulation were overrepresented compared to other groups. In contrast, sterilized soil at week 0 had a lower percentage of protein metabolism and respiration subsystems. Aside from the sterilized soil at week 0, the majority of metabolic pathways were similar between groups.

**Figure 4.**
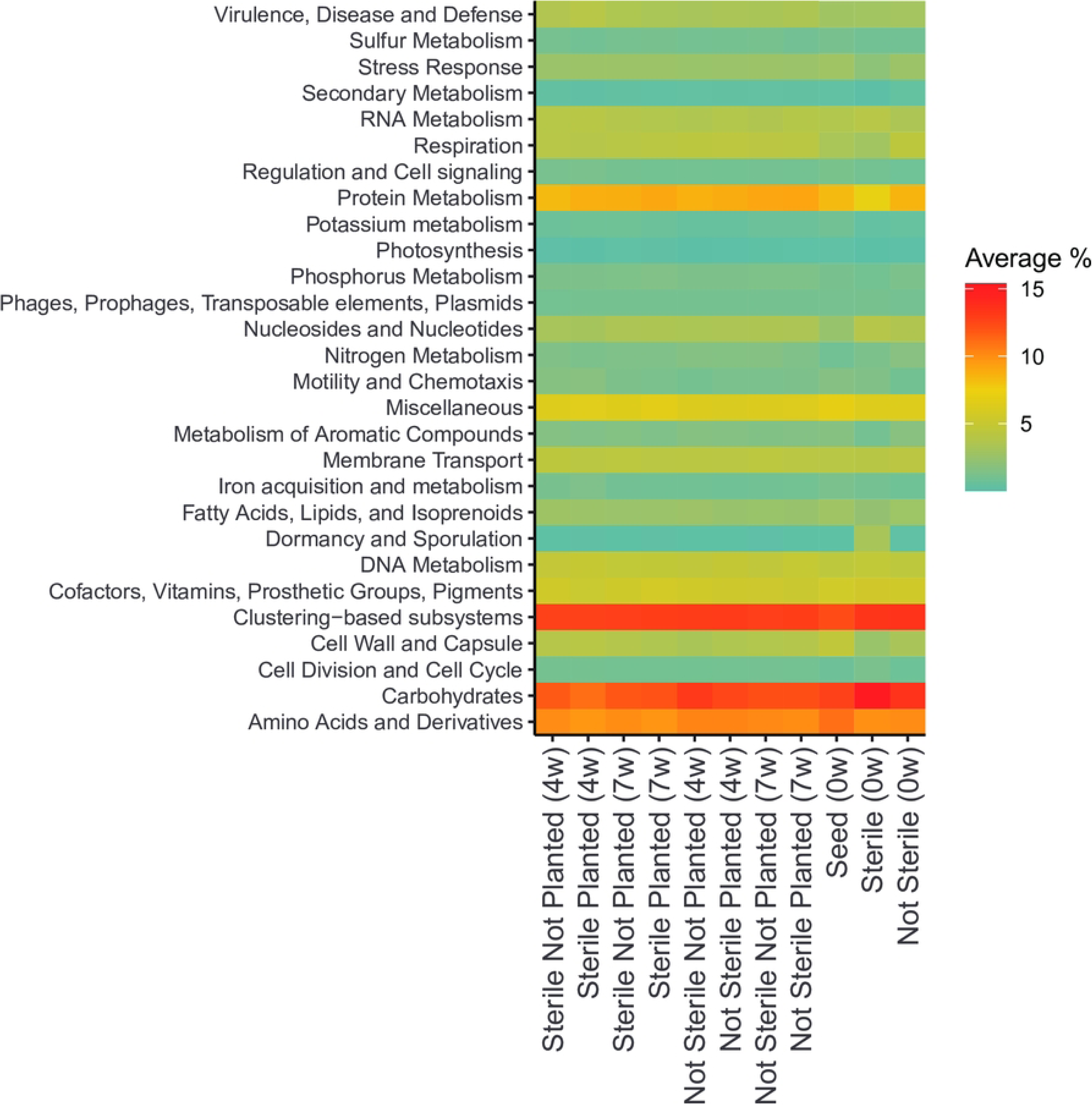
The impact of soil sterilization and planting with *Bouteloua curtipendula* on soil metabolic pathways. Potting soil was sterilized via autoclaving (as described in the materials and methods, “Sterile”) or left unsterilized (“Not Sterile”) and then added to 72 plug trays. *B. curtipendula* was seeded at week 0 (“Planted”) or left unseeded as a control (“Not Planted”). At four and 7 weeks, soil was collected for DNA purification and Illumina sequencing. Whole-genome sequencing was mapped to metabolic systems using MG-RAST. Colors represent the average percent of each metabolic system within a group (n=6; week 0 and seed, n=1).

Figure 5A shows the relative abundance (in percent) of selected level 3 subsystems across the treatment groups. Spore germination was overrepresented in sterilized soil at week 0 (Fig 5B), suggesting spore-forming bacteria were the primary species to survive autoclaving. Although underrepresented in sterile soil at week 0, Ton and Tol transport systems (within Membrane Transport) and iron acquisition were overrepresented in sterile planted soil at week 4 compared to all other groups (Fig 5D, I). Cobalt/Zinc/Cadmium resistance was also elevated in sterile planted soil at 4 weeks compared to other groups (Fig 5E). In contrast, nitrogen fixation and carbon monoxide-induced hydrogenase was underrepresented in sterile planted soil at 4 weeks compared to other groups (Fig 5G, H). By 7 weeks post-seeding, relative abundance of each subsystem in the sterile planted soil more closely resembled the other groups (Fig 5).

**Figure 5.**
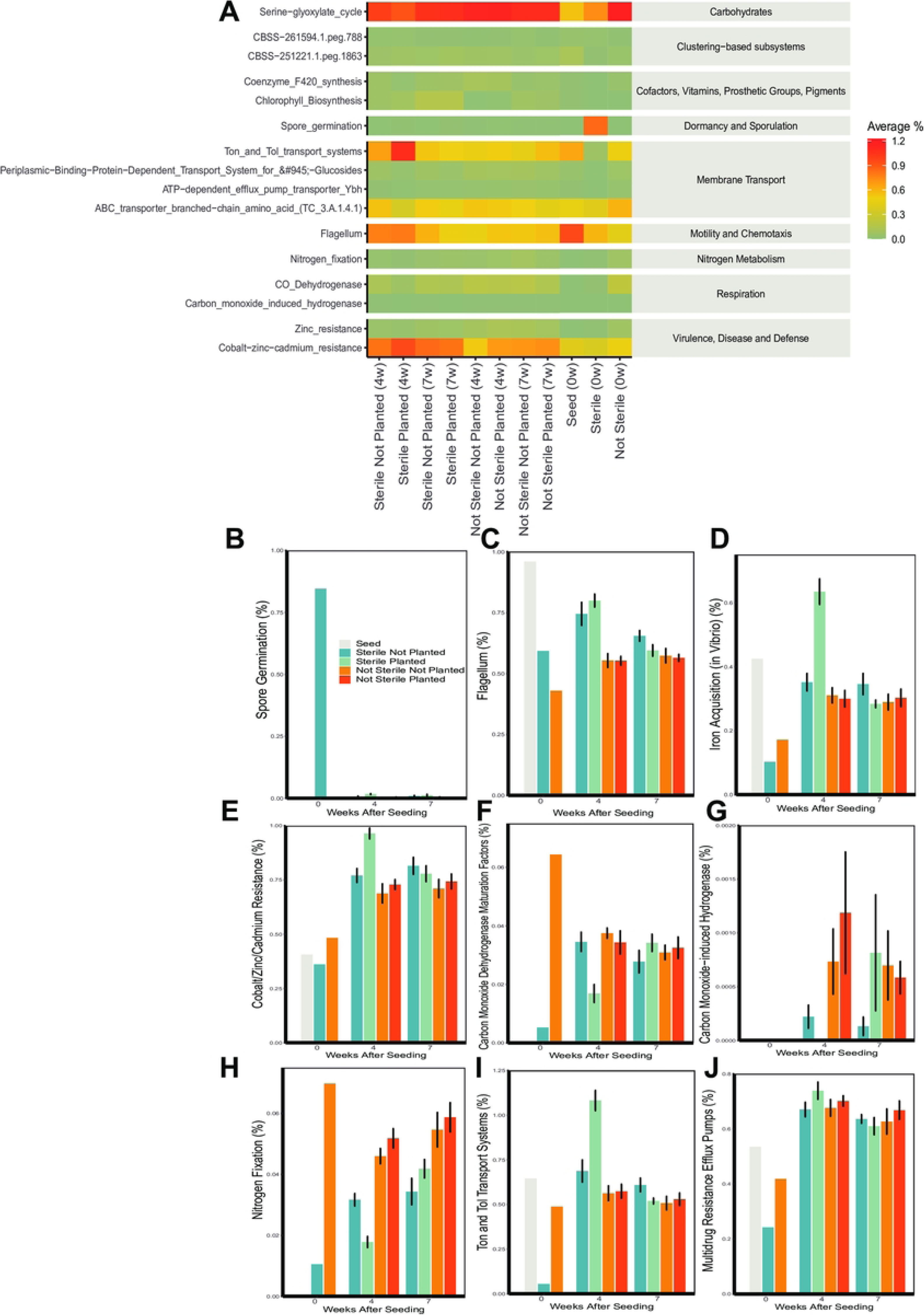
Soil sterilization and planting with *Bouteloua curtipendula* changed the relative abundance of soil metabolic pathway subsystems. Potting soil was sterilized via autoclaving (as described in the materials and methods, “Sterile”) or left unsterilized (“Not Sterile”) and then added to 72 plug trays. *B. curtipendula* was seeded at week 0 (“Planted”) or left unseeded as a control (“Not Planted”). At four and 7 weeks, soil was collected for DNA purification and illumine sequencing (as described in the materials and methods). Whole-genome sequences were mapped to level 3 functional systems in MG-RAST. (A) Select level 3 functional subsystems for each group with the indicated level 1 metabolic system (gray background, on right). (B) Mean +/- SEM of average percentage of select subsystems at weeks 0, 3, and 7 post-seeding (n=6; week 0/seed, n=1).

### Impacts of soil microbiome perturbation and planting with B. curtipendula on individual bacterial and fungal taxa

Almost all bacteria detected in the week 0 sterilized soil belonged to the phylum Firmicutes (Fig 6A). In contrast, not sterilized soil at week 0 was primarily composed of the phyla Proteobacteria, Planctomycetes, Bacteroidetes, and Actinobacteria (Fig 6A, Fig 7A, B). At 4 weeks post-seeding in the sterilized soil, Proteobacteria and Bacteroidetes were overrepresented compared to planted not sterilized soil (Fig 6A, Fig 7B). The phylum Actinobacteria was less abundant in sterilized planted soil at 4 weeks compared to not sterilized planted soil (Fig 6A, Fig 7A). By 7 weeks post-seeding, the relative abundance of Bacteroidetes, Actinobacteria, and Proteobacteria were similar between all groups (Fig 6A, Fig 7A, B).

**Figure 6.**
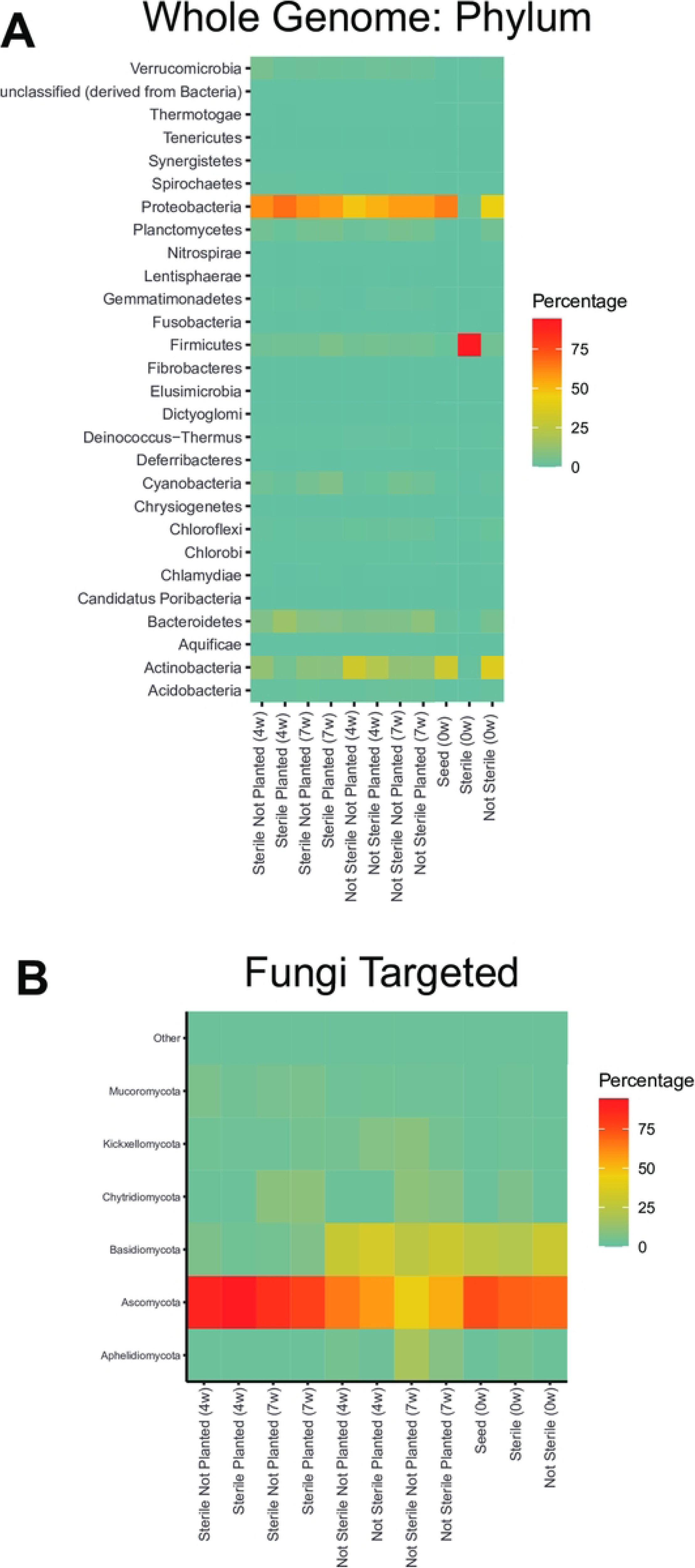
Soil sterilization and planting *Bouteloua curtipendula* alters the relative abundance of soil bacterial and fungal phyla. Potting soil was sterilized via autoclaving (as described in the materials and methods, “Sterile”) or left unsterilized (“Not Sterile”) and then added to 72 plug trays. *B. curtipendula* was seeded at week 0 (“Planted”) or left unseeded as a control (“Not Planted”). At four and 7 weeks, soil was collected for DNA purification and Illumina sequencing. Whole-genome sequences (A) or fungus ITS targeted sequences (B) of phyla representing greater than the threshold of 2% of the total rarefied reads are represented. Phyla below the threshold were pooled into “other.” Colors indicate the average percentage each phylum makes up within a group (n=6; week 0/seed, n=1).

**Figure 7.**
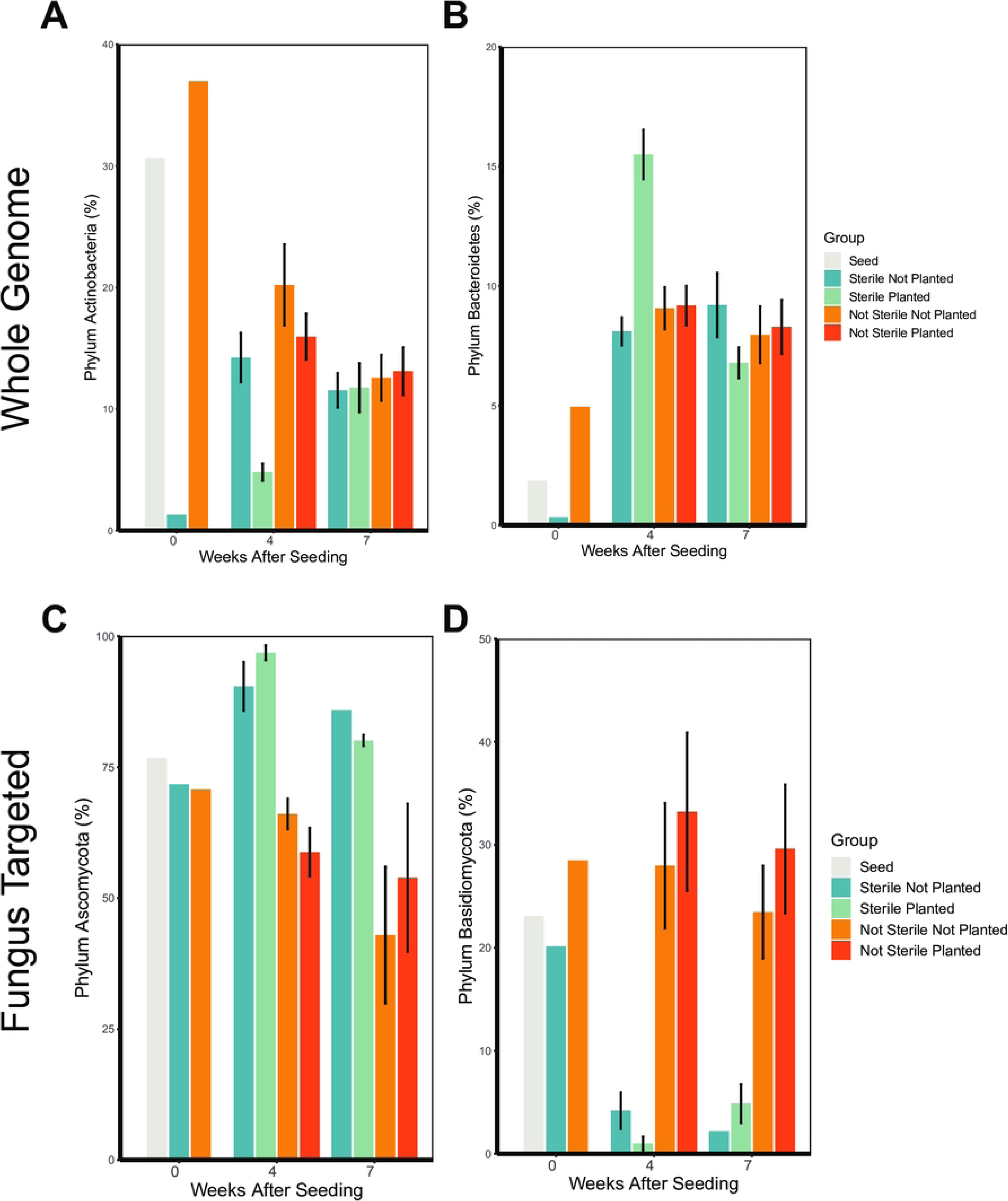
Soil sterilization and planting with *Bouteloua curtipendula* changes the relative abundance of bacterial phyla Actinobacteria and Bacteroidetes and fungal phyla Ascomycota and Basidiomycota. bacterial and fungal phyla. Potting soil was sterilized via autoclaving (as described in the materials and methods, “Sterile”) or left unsterilized (“Not Sterile”) and then added to 72 plug trays. *B. curtipendula* was seeded at week 0 (“Planted”) or left unseeded as a control (“Not Planted”). At four and 7 weeks, soil was collected for DNA purification and Illumina sequencing. Bars represent mean +/- SEM of bacterial phyla (A-B) Actinobacteria (A) and Bacteroidetes (B) or fungal phyla (C-D) Ascomycota (C) and (Basidiomycota) (D) (n=6; week 0/seed, n=1).

Amongst the fungi, Ascomycota, Basidiomycota, and Chytridiomycota were the most abundant in the soil (Fig 6B). Nearly all fungi on the seed belonged to the phyla Ascomycota or Basidiomycota, with an approximate 2.8:1 ratio of Ascomycota to Basidiomycota (Fig 6B, 7C-D). The fungal phylum Ascomycota was overrepresented in sterilized soil compared to not sterilized soil at both 4 and 7 weeks, representing greater than 75% of all fungal phyla reads (Fig 6B, Fig 7C). In contrast, the phylum Basidiomycota was less abundant in sterilized soils compared to unsterilized soils (Fig 6B, Fig 7D). Planting sterilized soil with *B. curtipendula* caused a slight increase in the relative abundance of Ascomycota and a slight decrease in the relative abundance of Basidiomycota at 4 weeks. By 7 weeks, planted soils showed slightly less Ascomycota and slightly more Basidiomycota. Interestingly, planted soils at week 4 had an approximate Ascomycota to Basidiomycota ratio of 30:1. By week 7, the Ascomycota to Basidiomycota ratio decreased to approximately 15.4:1, closer to the seed ratio of 2.8:1. These findings are consistent with the NMDS analysis showing the soil fungal community of sterilized soil planted with *B. curtipendula* resembles the seed more closely at 7 weeks post-seeding compared to 4 weeks post-seeding (Fig 2B).

For Bacteria, at the genus level, the sterilized soil at week 0 was almost entirely comprised of *Anoxybacillus*, *Bacillus*, *Geobacillus*, and *Paenibacillus* (Fig 8). These are all known to form heat-resistant spores and therefore not surprising to be found after sterilization. In contrast, the dominant genera in the not sterile soil at week 0 were *Mycobacterium*, *Streptomyces*, *Rhodopseudomonas*, *Salinispora*, *Mesorhizobium*, and *Bradyrhizobium* (Fig 8).

**Figure 8.**
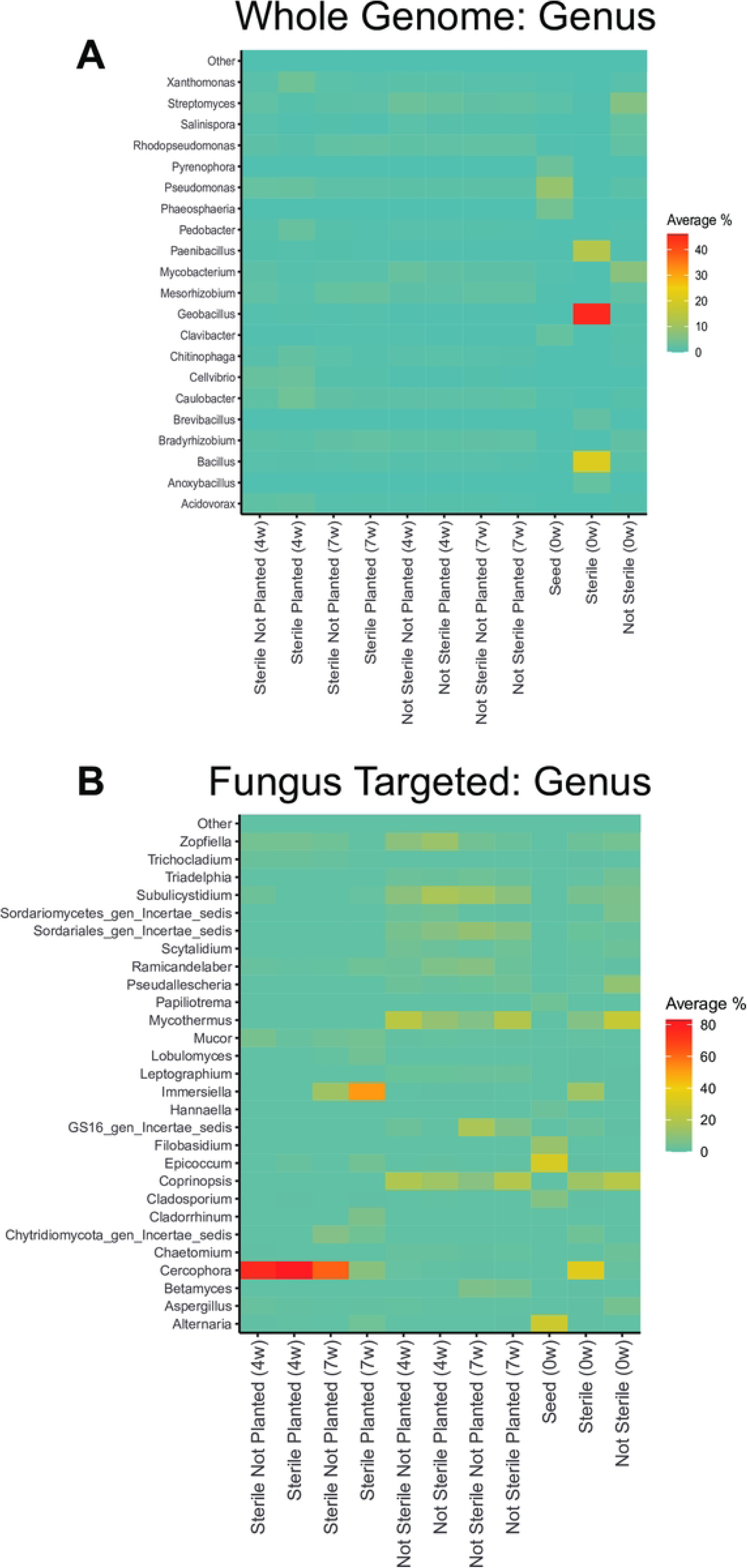
Soil sterilization and planting with *Bouteloua curtipendula* alters the relative abundance of soil bacterial and fungal genera. Potting soil was sterilized via autoclaving (as described in the materials and methods, “Sterile”) or left unsterilized (“Not Sterile”) and then added to 72 plug trays. *B. curtipendula* was seeded at week 0 (“Planted”) or left unseeded as a control (“Not Planted”). At four and 7 weeks, soil was collected for DNA purification and Illumina sequencing. Whole-genome sequences (A) or fungus ITS targeted sequences (B) of genera representing greater than the threshold of 2% of the total rarefied reads are represented. Phyla below the threshold were pooled into “other.” Colors indicate the average percentage each genus makes up within a group (n=6; week 0/seed, n=1).

Soil sterilization impacted the relative abundance of several bacterial genera at 4 weeks post-seeding. The genus *Caulobacter*, which is often associated with plants [42], was only induced by *B. curtipendula* seeded into sterile soil (Fig 9B). In addition, the plant pathogens *Acidovorax*, *Cellvibrio*, *Pseudomonas*, and *Xanthomonas* were all overrepresented in sterilized soils planted with *B. curtipendula* (Fig 9A-D). In contrast, *Rhodopseudomonas*, which is often categorized as a plant growth promoting bacteria, was significantly reduced in the sterilized soils planted with *B. curtipendula* compared to the not sterilized planted soils (Fig 9E). However, by week 7, *Rhodopseudomonas* relative abundance greatly increased the sterilized soils planted with *B. curtipendula* and *Rhodopseudomonas* was similarly abundant across all groups. Likewise, the potential plant pathogens *Acidovorax*, *Cellvibrio*, and *Xanthomonas* were no longer elevated in the sterilized soils by week 7 (Fig 9A-D).

**Figure 9.**
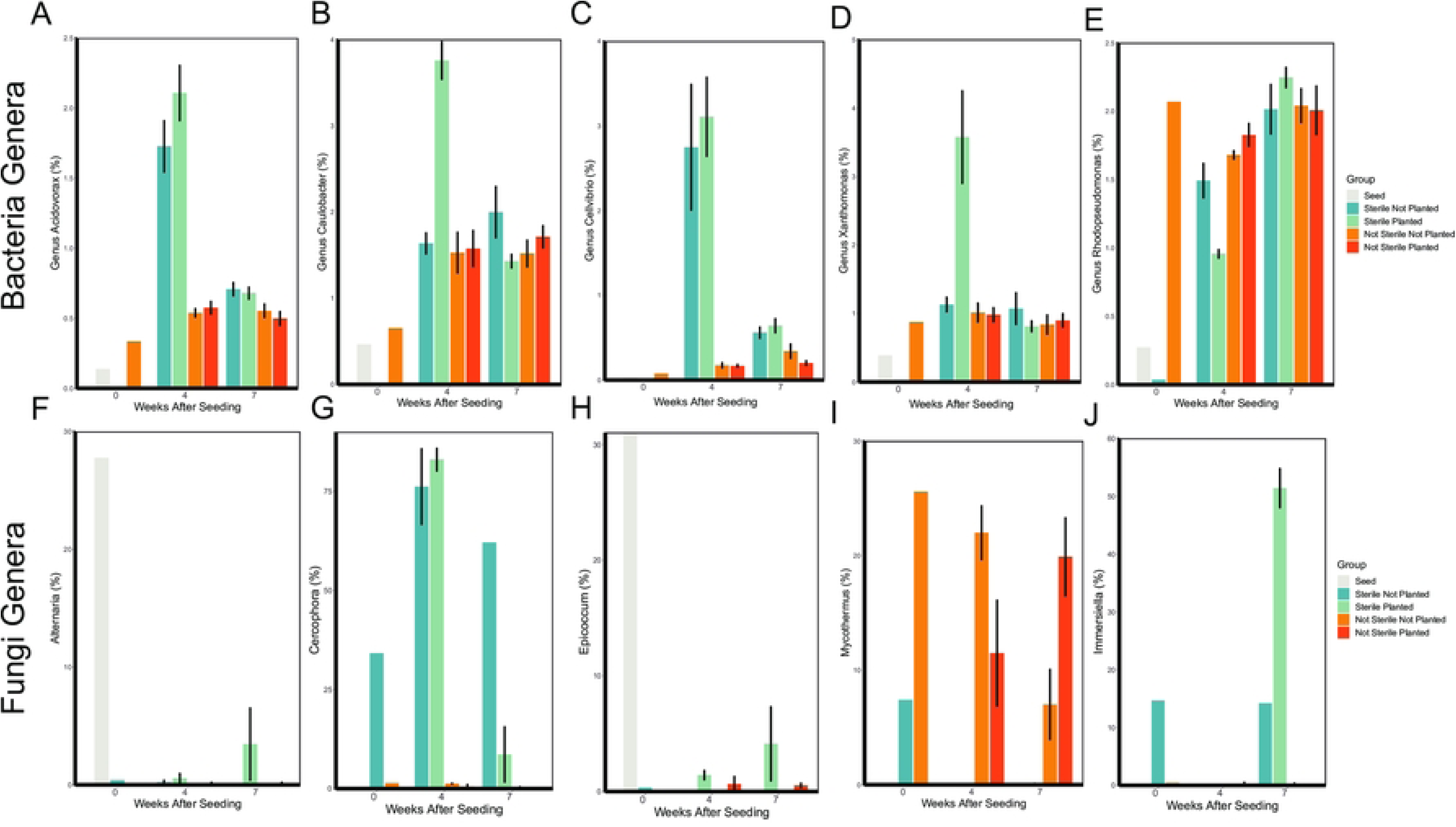
Soil sterilization and planting with *Bouteloua curtipendula* alters the relative abundance of specific bacterial and fungal genera. Potting soil was sterilized via autoclaving (as described in the materials and methods, “Sterile”) or left unsterilized (“Not Sterile”) and then added to 72 plug trays. *B. curtipendula* was seeded at week 0 (“Planted”) or left unseeded as a control (“Not Planted”). At four and 7 weeks, soil was collected for DNA purification and Illumina sequencing. Bars represent mean +/- SEM of bacterial (A-E) or fungal (F-J) genera (n=6; week 0/seed, n=1).

*Cercophora* was the predominant fungal genus in the sterilized soil at week 0 (Fig 8B, Fig 9G), while *Mycothermus* was the predominant fungal genus in the not sterilized soil at week 0 (Fig 8B, Fig 9I). *Epicoccum* and *Alternaria* were the most abundant fungal genera on the seed at week 0 (Fig 8B, Fig 9F, H). *Cercophora* continued to be the dominant fungal genus in the sterilized soil at 4 weeks in both the planted and unplanted conditions (Fig 8B, Fig 9G). By 7 weeks in sterilized planted soils, a decrease in *Cercophora* relative abundance coincided with an increase in *Epicoccum, Alternaria,* and *Immersiella*. In the sterilized soils not planted with *B. curtipendula*, *Cercophora* continued to be highly abundant at 7 weeks, with no apparent increase in *Epicoccum*, *Alternaria*, or *Immersiella*. Interestingly, *Cercophora*, *Immersiella*, and *Epicoccum* remained low in abundance in the not sterile soils throughout the experiment (Fig 8B, Fig 9G, H). Instead, *Mycothermus* remained the predominant fungal genus in the not sterile soil through week 7 (Fig 8B, Fig 9I).

However, while *Mycothermus* decreased in relative abundance over time in the not planted soil, the genus increased in relative abundance from 4 weeks to 7 weeks in planted soil (Fig 9I). The observation that seed-associated fungi (*Alternaria* and *Epicoccum*) are good at colonizing the sterile soil, but not the not sterile soil is consistent with the earlier NMDS overall community observations.

## Discussion

In this study, we show how transient perturbation of the soil microbial community through autoclave sterilization impairs the growth of the prairie grass *B. curtipendula. B. curtipendula* seeded into sterilized soil had a slower germination, fewer grass blades and shorter blade length throughout the course of a 7-week controlled experiment, but the difference was most notable in the first 4 weeks post-seeding. Although sterilization was transient, the soil microbial community did not immediately return to a pre- sterilization state. Instead, planting *B. curtipendula* seed into the sterilized soil caused a strong bacterial community divergence at 4 weeks that eventually stabilized by 7 weeks post-seeding (Fig 2A). In contrast, the fungal community divergence in the sterilized soil planted with *B. curtipendula* was slower, not being detectable until 7 weeks post- seeding (Fig 2B). *B. curtipendula* seeded into sterilized soil also induced more of the potentially pathogenic bacterial genera *Acidovorax*, *Cellvibrio*, and *Xanthomonas* and less of the plant growth promoting *Rhodopseudomonas* at 4 weeks post-seeding (Fig 9A-E). This overabundance of bacterial plant pathogens subsided at 7 weeks post- seeding into sterilized soil, giving way to *B. curtipendula*-induced fungal species such as *Alternaria*, *Epiccocum*, and *Immersiella* (Fig 9F,H, I).

Although two rounds of autoclaving did not completely sterilize the soil, it did greatly reduce bacterial richness and diversity (Fig 5A, C). The remaining bacterial taxa identified through high throughput sequencing were spore forming bacilli (Fig 8A). Also, the dominant functional subsystem in sterilized soil at time 0 was spore germination (Fig 4A, B). The genetic data are consistent with the morphology of viable bacteria that grew on nutrient agar plates following our sterilization protocols (data not shown). Fungal richness and diversity were only moderately reduced immediately following soil sterilization (Fig 5B, D). However, the low diversity and richness of fungi genera at 4 weeks post-seeding suggests the sequences detected immediately after sterilization arose from non-viable fungi. Indeed, we did not observe significant fungal growth on nutrient agar plates of sterilized soil even after one week of culturing at 20 degrees C (data not shown). Thus, the sterilization caused a significant disruption to the soil microbial communities, where the bacteria show a more acute stress response and faster recovery than fungi in the plant soil microbiome.

Our results demonstrate soil microbial communities are highly dynamic and influenced by *B. curtipendula* planting following acute destabilization. While the overall bacterial community in sterilized soil was distinct from that of not sterilized soil at 4 weeks (Fig 2A), this difference was exacerbated when sterilized soils were planted by *B. curtipendula*. In sterilized soils, *B. curtipendula* significantly decreased the relative abundance of the phylum Actinobacteria and increased the relative abundance of the phylum Bacteroidetes at 4 weeks post-seeding. In not sterilized soils, the bacterial community was stable and not significantly impacted by seeding *B. curtipendula*.

We also report how *B. curtipendula* showed delayed signs of germination (Fig 1A) and grew more slowly (Fig 1B) in soils with an acutely disrupted microbiome compared to unperturbed soils (Fig 1). It is interesting to note how soil sterilization increased the relative abundance of potential plant pathogen bacteria, such as *Acidovorax* [43–45], *Cellvibrio* [46,47], and *Xanthomonas* [48,49] at 4 weeks post-seeding. The latter observation is consistent with reports of *Xanthomonas* increase in phytopathogenic biofilm formation in plant roots [50,51]. By 7 weeks post-seeding, the potentially pathogenic bacteria stabilized and returned to levels comparable to that of the not sterilized soil seeded with *B. curtipendula*. These data are consistent with our observation of impaired *B. curtipendula* growth in sterilized soil, especially during the first 4 weeks post-seeding.

Acute disruption of gut microbiota with antibiotics is known to increase the relative abundance of human pathogens [36]. Our findings suggest acute disruption of the soil microbiota could also promote plant pathogens that impede growth of plants, such as the prairie grass *B. curtipendula*. It is interesting that the bacterial communities restabilized by 7 weeks post-seeding and *B. curtipendula* growth in sterilized soil was less deficient at these later time points. It is possible that soil texture, moisture, pH, and organic mineral content (which should be largely similar between sterilized and not sterilized soil) is the major determinant of long-term soil microbial community, and acute disturbance will always be a transient disruption before returning to the long-term stable community. It is also possible that the fungal species aided in stabilization of the bacterial community and the reduction of potentially pathogenic bacterial taxa by 7 weeks. Indeed, we find the *B. curtipendula*-induced fungal community took 7 weeks to establish in sterilized soil (Fig 2B). Interestingly, one of the fungal genera that increased in *B. curtipendula*-planted sterilized soil by 7 weeks include*s Epicoccum*, which was also highly abundant on the *B. curtipendula* seed. Thus, seed-associated fungi could be responsible for reducing the plant pathogens in the soil and thus promoting *B. curtipendula* growth. Indeed, *Epicoccum sp.* associated with other plants have been shown to produce antimicrobial compounds against phytopathogens [52]. In contrast, the other seed-associated fungi induced by 7 weeks, *Alternaria*, is most commonly associated as a fungal plant pathogen itself [53,54]. Future studies will be necessary to determine whether seed-associated fungi promote *B. curtipendula* growth through antimicrobial activity against plant pathogens in the soil. Nevertheless, our results show the soil microbial dynamics associated with inhibition of prairie grass growth following acute soil sterilization.

Our study is consistent with previous studies showing interactions between plants and the soil microbial community. Much of this work involves a group of bacteria categorized as plant growth promoting bacteria (PGPB) and fungi (PGPF) [55].

Microbes known to be PGPB and PGPF can release secondary metabolites that suppress plant pathogenic bacteria and fungi in the soil, thus promoting plant growth [56]. Such is the case for the fungus *Mycothermus*, which showed a strong positive correlation with Lisianthus (*Eustoma sp.*) growth due to suppression of plant disease [57]. PGPB can also promote nutrient uptake and increase fertilizer efficiency [58]. One specific bacteria genus associated with plant growth is *Rhodopseudomonas*. When inoculated into sterilized soil, *Rhodopseudomonas* enhanced germination of tomato plants [59]. *Rhodopseudomonas* incoculation also promoted growth of Stevia [60], Brassica plants [61,62], beans (*Vigna mungo*) [63], rice [64,65], and provide systemic resistance to TMV in tobacco plants [66]. We found significantly less *Rhodopseudomonas* in sterilized soil planted with *B. curtipendula* compared to not sterilized soil planted with *B. curtipendula* at four weeks post seeding (Fig 9). In addition, we found sterilization essentially eliminated *Mycothermus* from the soils throughout the course of our experiment (Fig 9). The reduction of *Rhodopseudomonas* and *Mycothermus* in sterilized soil is associated with increased pathogenic bacterial and fungal taxa (Fig 9) and impaired *B. curtipendula* growth (Fig 1). Thus, our results suggest that a diverse and unperturbed microbial community could support prairie grass growth through multiple mechanisms: directly promoting plant growth and reducing the abundance of potential plant pathogens.

It has been shown that the microbial interaction with plant roots can result in improved nutrient and mineral uptake, help in plant-growth promotion as well a suppression of phytopathogens [67,68]. The role of plant growth promotion involves phytohormone production, nitrogen fixation, siderophore production, solubilization of inorganic substances (P, K, Zn etc.), and the microbial remediation of heavy metal toxicity (such as high cadmium in fertilizer) [68,69]. The mechanisms of suppression of phytopathogens can involve direct methods of production of antibacterial or antifungal metabolites or production of wall-degrading enzymes such as chitinase, but also involves indirect routes such as iron chelation and depletion from the rhizosphere [70]. A search of the known genomes of *Rhodopseudomans* strains [71] shows that several strains contain genes for 1-aminocyclopropane-1-carboxylate (ACC) deaminase, which is an enzyme known to reduces plant stress hormones [28], and genes for enzymes involved in indole-3-acetic acid (IAA) and 5-aminolevulinic acid (ALA) production, which may also be one of the mechanisms of plant growth enhancement [59]. Certain strains of *Rhodopseudomonas* have been shown to have plant growth promoting effects by stimulating nitrogen uptake and increasing IAA levels [62], which is consistent with our observations here. Unfortunately the current resolution of metagenomic sequencing is unable to confidently identify to the species level, nevertheless it is likely that the *Rhodopseudomonas* strains involved here have one or several of the characteristics above that contribute to the plant growth promoting effect.

The subsystem ‘Zinc resistance’ is one of the Virulence, Disease, and Defense (Subsystem 1) functions, that was elevated in sterile planted soil at 4 weeks compared to other groups (Fig 5). The specific enzymes related to this (Function level) were the sensor protein and response regulator of the Sigma-54 two component system. The Sigma factor 54 is a central regulator in many pathogenic bacteria and has been linked to important functional traits such as motility, virulence, host colonization, and biofilm formation [72]. The higher level of this function in the sterile soil conditions is consistent with our observation of higher levels of potential pathogens in the sterile conditions, before a more diverse microbiome becomes established. Overall the majority of metabolic pathways were similar between groups and conditions, aside from the sterilized soil at week 0, which indicates an overall metabolic stability in the microbiome, where the major soil biochemical reactions can be performed by a diverse group of bacteria.

Ultimately, our results suggest that functional and diverse microbial soil communities could aid in the establishment of prairie grasses and subsequently increase the success of urban prairie plantings. Native plant species promote broad ecological functions in habitat areas and these functions are reduced by invasive species. However, microbial communities associated with invasive species can favor invasive grasses and impede native grasses [73]. Another study found soil sterilization increased the success of invasive *Bromus tectorum* infiltrating into *Bouteloua gracilis* (a plant closely related to *B. curtipendula* used in our study) [37]. These findings combined with the results of our study emphasize the importance of soil microbial communities in promoting highly functional native plant communities. Furthermore, our study indicates that the bacteria are involved in the initial establishment of beneficial conditions that paves the way for a solid fungal and plant seedling development. Understanding these microbiome-plant relationships in native *B. curtipentula* opens up possibilities for incorporating target strains that help the colonization and growth of this native grass in restored native habitats. One way to approach this might be to incorporate these growth-promoting strains into seeds, similar to the ENDOSEED concept that has been performed in crop plants [27,74]. Irrespective of the application method, efforts to promote native habitat spaces should consider approaches which include native-plant promoting soil microbial communities to ensure the greatest habitat functionality.

## Acknowledgements

This work was supported by the Wilson Enhancement Fund for Applied Research at Bellevue University. We would like to thank Julian Ramirez for his assistance in maintaining the greenhouse and plants used in this experiment.

